# Human Bone Marrow Assessment by Single Cell RNA Sequencing, Mass Cytometry and Flow Cytometry

**DOI:** 10.1101/416750

**Authors:** Karolyn A. Oetjen, Katherine E. Lindblad, Meghali Goswami, Gege Gui, Pradeep K. Dagur, Catherine Lai, Laura W. Dillon, J. Philip McCoy, Christopher S. Hourigan

**Author notes:** To whom correspondence should be addressed: Christopher S. Hourigan Laboratory of Myeloid Malignancies Hematology Branch National Heart, Lung and Blood Institute Room 10CRC 5-5130, 10 Center Drive Bethesda, Maryland, 20814-1476, USA.

## Abstract

New techniques for single-cell analysis have led to insights into hematopoiesis and the immune system, but the ability of these techniques to cross-validate and reproducibly identify the biological variation in diverse human samples is currently unproven. We therefore performed a comprehensive assessment of human bone marrow cells using both single-cell RNA sequencing and multiparameter flow cytometry from twenty healthy adult human donors across a broad age range. These data characterize variation between healthy donors as well as age-associated changes in cell population frequencies. Direct comparison of techniques revealed discrepancy in the quantification of T lymphocyte and natural killer cell populations. Orthogonal validation of immunophenotyping using mass cytometry demonstrated good correlation with flow cytometry. Technical replicates using single-cell RNA sequencing matched robustly, while biological replicates showed variation. Given the increasing use of single-cell technologies in translational research, this resource serves as an important reference dataset and highlights opportunities for further refinement.

## Introduction

New technologies for characterizing cell populations are being implemented to more deeply describe the cell surface receptor phenotype and gene transcriptional signature at the single cell level (1, 2). Benefits of single cell approaches include examination of heterogeneity within the sample, and the most recent advances permit use of samples with very limited cell numbers for high dimensional characterization of cell surface phenotype or transcriptome. Single cell RNA sequencing (scRNAseq) has been used to elucidate hematopoietic differentiation (3-5) and immune cell subsets (6) including dendritic cells and monocytes (7), and innate lymphoid cells (8). Mass cytometry has been applied to the study of tissue-infiltrating immune cells (e.g. melanoma (9), renal cell (10), lung (11), and breast (12) cancers).

Expanding these new single cell approaches to patient samples requires a clear understanding of their correlation with established techniques, including flow cytometry. In order to facilitate and validate analysis of large databases of scRNAseq we set out to provide a data set of human bone marrow analyzed by both scRNAseq and deep immunophenotyping. Our reference cohort includes a broad range of donor ages in recognition of age-related variation in the healthy population.

## Materials and Methods

### Bone Marrow Aspirate Collection

Healthy volunteers were recruited for bone marrow aspirate collection at the National Institutes of Health. This research was approved by the National Heart, Lung and Blood Institute Institutional Review Board, and all participants provided oral and written informed consent. Using standard operating procedures, mononuclear cells from bone marrow aspirates were isolated using Ficoll density gradient separation and cryopreserved in 90% FBS/ 10% DMSO for storage in liquid nitrogen. Assays were performed as listed in Table 1 using matched cryopreserved vials from each donor.

**Table 1.**
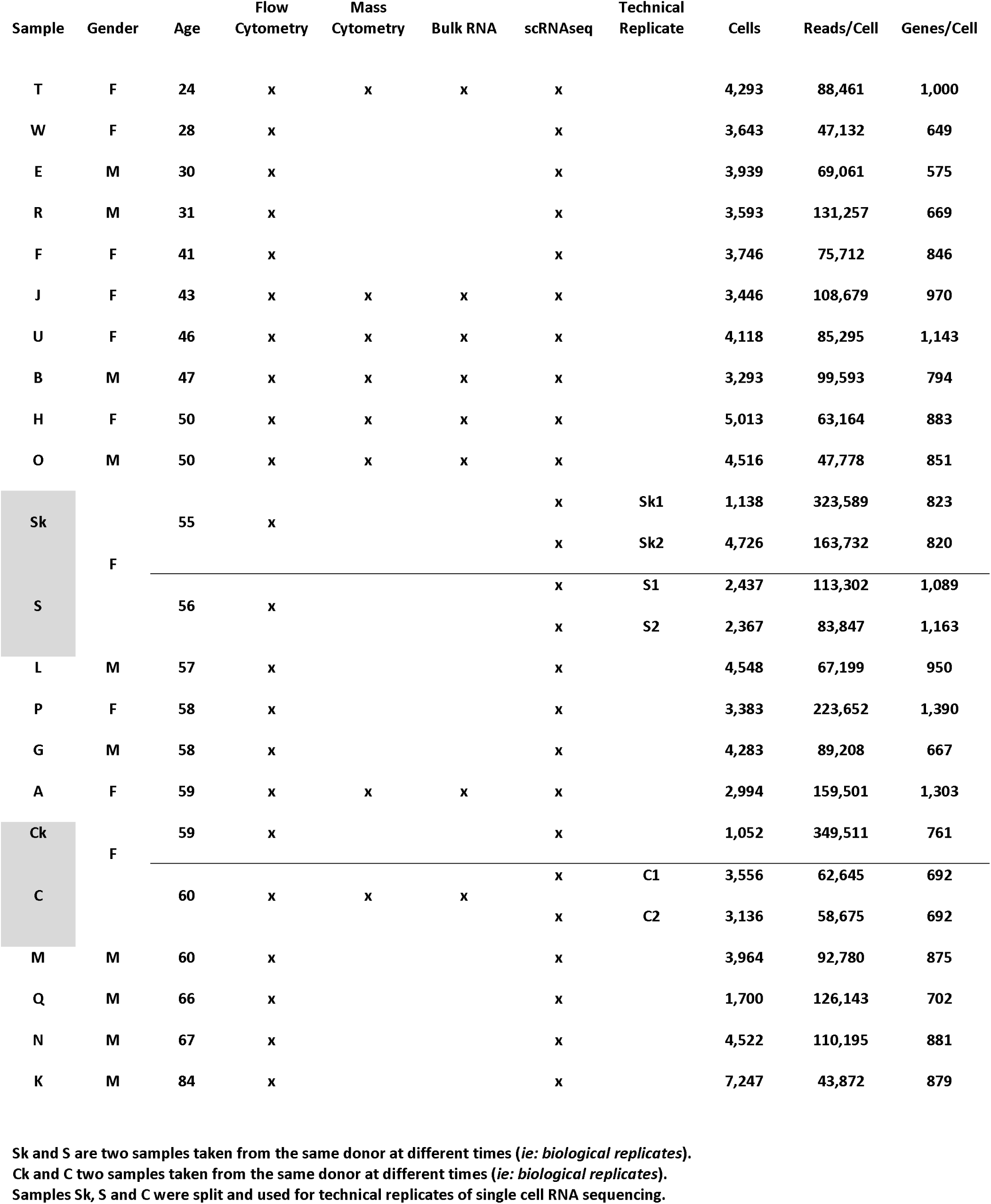
Heathy volunteer sex and age at time of bone marrow aspiration. Biological replicate time points for a second longitudinal bone marrow aspirate from the same volunteers are shown within grey boxes. Assays from matched cryopreserved bone marrow mononuclear cell vials are indicated. Single cell RNA sequencing cell counts and sequencing depth (reads per cell and genes per cell) are listed for each donor and replicate.

### Single cell RNA Sequencing

scRNAseq was performed using 10X Genomics Single Cell 3’ Solution, version 2 according to manufacturer’s instructions (protocol rev A). Libraries were sequenced on HiSeq3000 and analyzed using Cell Ranger V2.0.0 (10X Genomics). Quality control metrics were used to select cells with mitochondrial gene percentage less than 8% and at least 500 genes detected. Samples were analyzed using Seurat (www.satijalab.org/seurat) using canonical correlation analysis with Louvain clustering, and visualized by t-distributed stochastic neighbor embedding (tSNE) (31). Developmental trajectories were created using Monocle versions 2 and 3 (32-34), the latter using Uniform Manifold Approximation and Projection for Dimension Reduction (UMAP) (35).

### Flow cytometry

BMMCs were thawed in RPMI-1640 (Gibco) with 10% FBS and resuspended in cell staining buffer. Benzonase nuclease (Sigma Aldrich, catalog #E1014-25KU) was added for some samples during thawing to minimize cell clumping. Cells were blocked with Human TruStain FcX Fc receptor blocking solution (Biolegend, catalog #422302) and stained with antibodies listed in Table S1 followed by LIVE/DEAD Fixable Yellow stain (Life Technologies Corporation, Grand Island, NY) and fixation with 1% formaldehyde. Data were acquired with a Becton–Dickinson LSRFortessa (BD, San Jose, CA, USA) equipped with five lasers (355, 407, 488, 532 and 633 nm wavelengths) and 22 PMT detectors using DIVA 8 software using the high throughput sampler (BD) system at a flow rate of 2.5ul/sec in a 96 well U bottom tissue culture plate. Compensation controls were performed using single color staining of compensation beads (BD), and daily quality assurance was performed using Cytometer setup and Tracking beads (BD) as per manufacturer’s recommendation along with 1 peak rainbow Beads (BD) and 8 peak beads (Spherotec)(36, 37). Post-acquisition analysis was performed using Flowjo 9.9.6 (Treestar Inc., San Carlos, CA, USA). Analysis excluded debris and doublets using light scatter measurements, and dead cells by live/dead stain. Gating strategies used to identify immune cell subsets are provided in Figure S2.

### Mass cytometry

Thawed BMMCs were stained for 37 markers using the MaxPar Complete Human T Cell Immuno-Oncology Panel Set (Fluidigm), according to manufacturer instructions. Briefly, cells were thawed, washed, incubated with cisplatin cocktail for viability, fixed in 1.6% formaldehyde and permeabilized. Cells were then stained with the antibody cocktail, incubated with intercalation solution, mixed with EQ4 element beads and acquired with a Helios mass cytometer (Fluidigm). Gating and viSNE analysis (38) were performed using Cytobank (cytobank.org). Initial analysis excluded doublets using DNA content and non-viable cells using cisplatin. CD45-positive cells were gated for viSNE analysis of 100,000 total events from all analyzed samples.

### Bulk RNA sequencing

RNA was harvested from thawed cell vials of BMMCs using AllPrep kits (QIAGEN). Libraries were prepared using TruSeq Stranded Total RNA Sample Preparation Kit (Illumina) with 1ug of RNA input. Sequencing was performed by paired-end 75 nt on Illumina HiSeq 3000. Fastq files were mapped to using *kallisto*, and gene counts were tabulated using *tximport*. Deconvolution was performed using Xcell v1.1 (xcell.ucsf.edu)(16) or Cibersort using LM22 gene signature and 100 permutations (cibersort.stanford.edu)(17).

### Data analysis and statistics

Data were analysis, visualization and statistical comparisons were performed R (cran.r-project.org). Bland-Altman analysis (39) was implemented in the BlandAltmanLeh package v0.3.1.

### Data availability

FCS files for flow cytometry and mass cytometry data sets have been deposited in FlowRepository (*accession#*). Single cell RNA sequencing and bulk RNA sequencing datasets have been deposited in Gene Expression Omnibus (GEO) (*accession#*).

## Results

### Healthy donor characteristics

Twenty healthy volunteers were recruited for bone marrow aspiration procedures. The cohort consisted of 10 males and 10 females with ages ranging 24-84 years old and median age of 57 years. A second bone marrow aspiration was performed for two donors (Ck, Sk) (*“biological replicate”*) either 2 or 5 months after their first aspiration respectively. Cryopreserved cells from all twenty donors were analyzed by droplet-based scRNAseq and flow cytometry, and additional cryopreserved vials for eight donors were analyzed by mass cytometry for T cell phenotyping, as well as bulk RNA sequencing, as summarized in Table 1.

### Single cell RNA sequencing

Droplet-based scRNAseq of bone marrow mononuclear cells (BMMCs) for all donor samples was performed with goal minimum sequencing depth of 50,000 reads/cell and detected a mean of 880 genes/cells (range 575-1,390 gene/cell, Table 1). Greater than 90,000 cells were captured; using quality filters of at least 500 genes per cell and less than 8% mitochondrial RNA content, 76,645 cells were analyzed in the final analysis.

To account for sample variations between donors, alignment of all samples was performed in Seurat using canonical correlation analysis (CCA) then visualized using t-distributed stochastic neighbor embedding (t-SNE). Cell clusters were distinguished using the Louvain clustering algorithm implemented in Seurat. Compiled analysis of all donor cells is annotated in Figure 1A with the contribution of each individual donor displayed in Figure S1. All major previously identified populations of bone marrow mononuclear cells were present in the clustered scRNAseq analysis.

**Figure 1.**
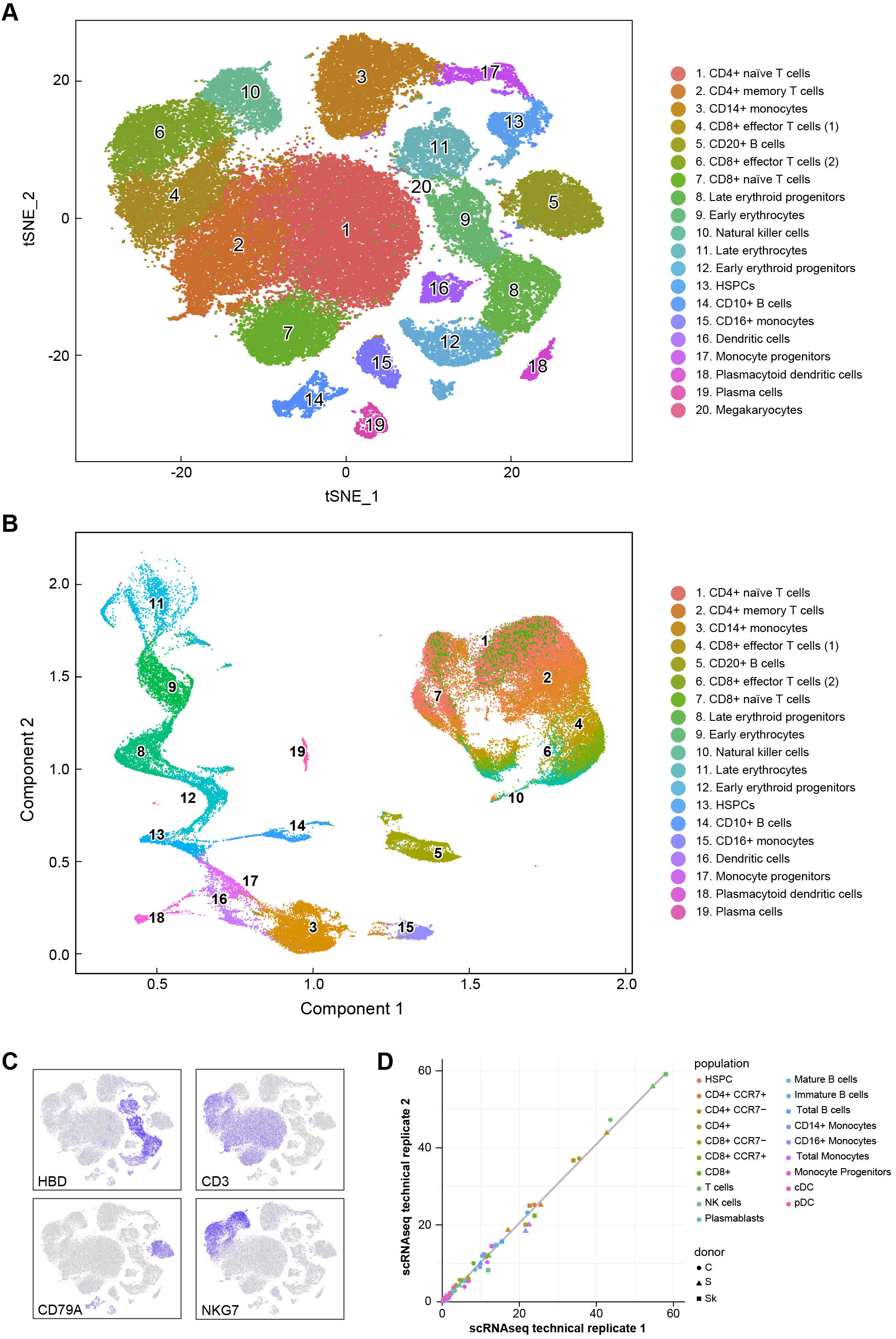
Single cell RNA sequencing of healthy bone marrow cells. (A) Cluster identification visualized using t-distributed stochastic neighbor embedding (t-SNE). (B) Single cell trajectory analysis using UMAP/Monocle 3. Color as in Figure 1A. (C) Examples of canonical gene expression used for annotation. (D) Reproducibility of technical replicates for single cell RNA sequencing. Linear regression line displayed in grey.

Single cell trajectory analysis was performed using Monocle 3. As there were potentially multiple disjoint trajectories in this complex dataset containing a large number of cells, UMAP was used for dimension reduction. The resulting development trajectories clearly display the major lymphoid, myeloid and erythroid lineages of hematopoiesis with correct ordering of developmental stages (Figure 1B). Trajectories of erythroid and myeloid lineages could also be created using an earlier, well validated, version of this software (Monocle2, see Figure S1) and were consistent with those observed for the full dataset.

Annotation of cell cluster identities was determined using a panel of canonical gene expression, with the expression patterns for a subset of these genes displayed in Figure 1C. Analysis of each donor sample individually using principal component analysis (PCA) in Seurat revealed suboptimal quantification of frequencies of some transcriptionally similar cell subsets, including those annotated as effector T cells and NK cells. Such clusters were typically well delineated for each individual sample when using CCA in the context of the entire dataset (Figure S1).

A potential use of scRNAseq is to compare across two or more samples. To confirm the validity of scRNAseq for this approach, assay reproducibility was determined by preparing duplicate, side-by-side libraries from cells thawed from the same cryopreserved vial, for a total of three cryopreserved samples. Cell subtype quantification for each of these technical replicate pairs matched robustly (Figure 1D). The optimum number of cells required to identify, using scRNAseq, sub-populations within a heterogenous samples remains an area of interest (13). Technical replicates ranged from 1,138 to 6,692 cells from the same sample (Table 1).

### Flow cytometry

13-color flow cytometry using five customized panels (“T, B, NK, Mono and DC”, see Table S1) designed to allow deep immunophenotyping of the predominant cell populations found in human bone marrow was performed on all samples. Approximately 1 million cells were stained for each panel, and a median of 196,000 CD45 positive events collected (25^th^-75^th^ percentile: 100,000-278,000 events). Gating strategy is shown in Table S2. Most frequent cell subtype populations observed were, in order, T cells, monocytes, B cells, natural killer cells (NK), dendritic cells (DC) and hematopoietic stem/progenitor cells (HSPC) (see Figure S2).

Paired analysis of the same sample by both transcriptome and cell surface phenotype offers a powerful opportunity to compare cell population frequencies determined by these methods. The proportion of major cell populations is summarized for scRNAseq and flow cytometry in Figure 2A. Sample-by-sample correlations for all of these populations are shown in Figure 2B. It is well established that the T memory cell population increases with increasing age in humans, likely due to response to viral infection (in particular CMV), and this trend was reproduced in our cohort using both scRNAseq and flow cytometry (Figure S2)(14). Two subjects had a second bone marrow aspiration performed at either 2 or 5 months after their first aspiration. These biological replicates showed good concordance by flow cytometry but showed variation by scRNAseq particularly in lower frequency cell subsets, likely from sampling error (Figure S2)

**Figure 2.**
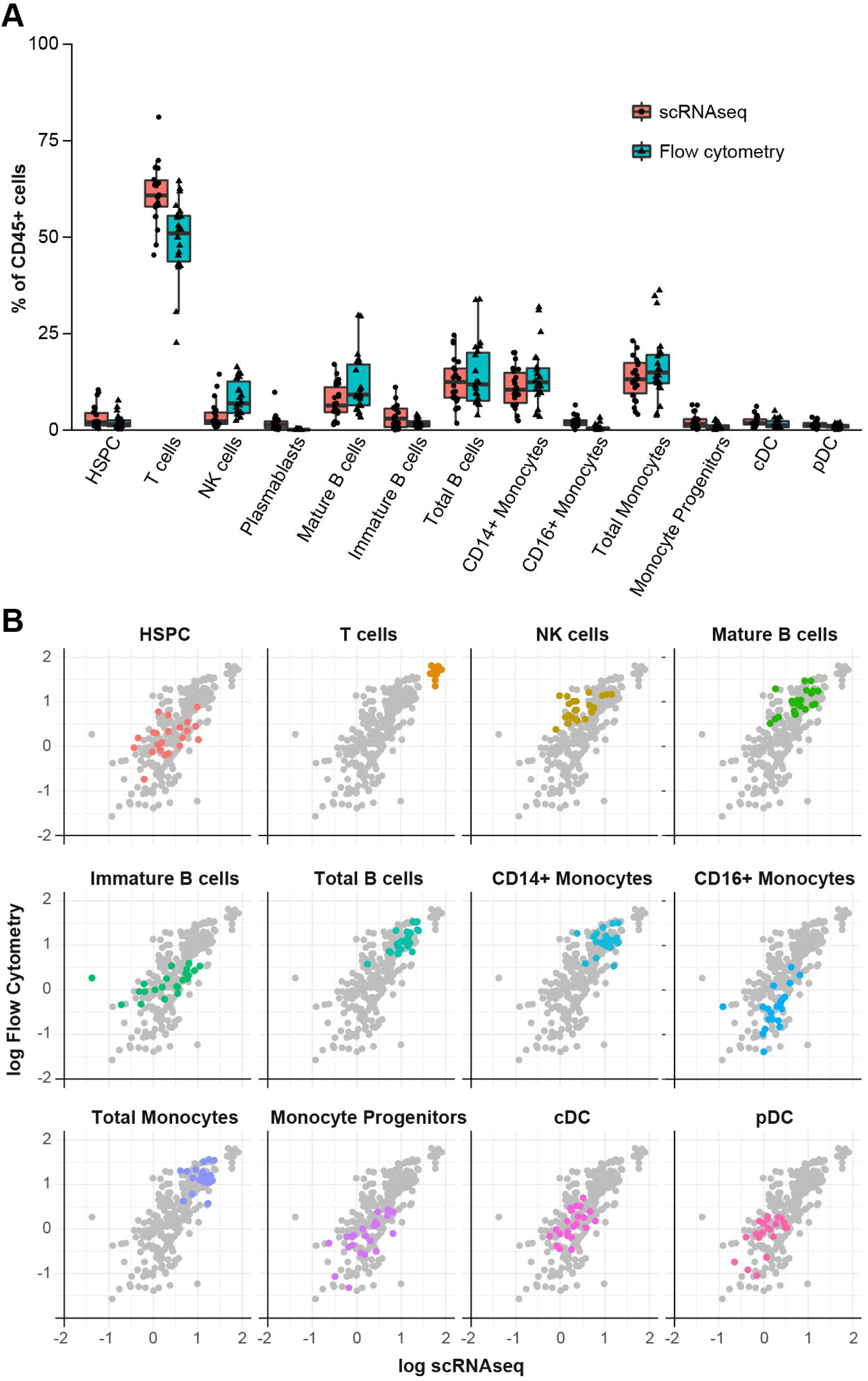
Comparison of single cell RNA sequencing and flow cytometry assessment of bone marrow cell type population frequencies. (A) Frequencies for major cell populations in human bone marrow shown for single cell RNA sequencing and flow cytometry. (B) Individual sample comparisons by scatter plot for each cell population. All population comparisons are shown in background in grey. Population frequencies are reported using denominator of all CD45 positive cells.

While concordance between these two modalities was generally good, it appeared that T cell frequency was elevated, and NK cell frequency decreased in scRNAseq as compared with flow cytometry. This led to a more detailed examination of T cells subsets and orthogonal validation of cell surface immunophenotyping using a third single cell modality.

### Mass cytometry

In order to more deeply characterize immune populations within healthy bone marrow, and to validate our flow cytometry results, T cell phenotyping was performed by mass cytometry using a 37-marker panel for a subset of eight donors. Using Cytobank software, CD45-positive cells were visualized using viSNE across the panel of markers (Figures 3A and S3). Correlation between mass cytometry and flow cytometry for CD4- and CD8-positive T lymphocyte subsets was good as shown in Figure 3B.

**Figure 3.**
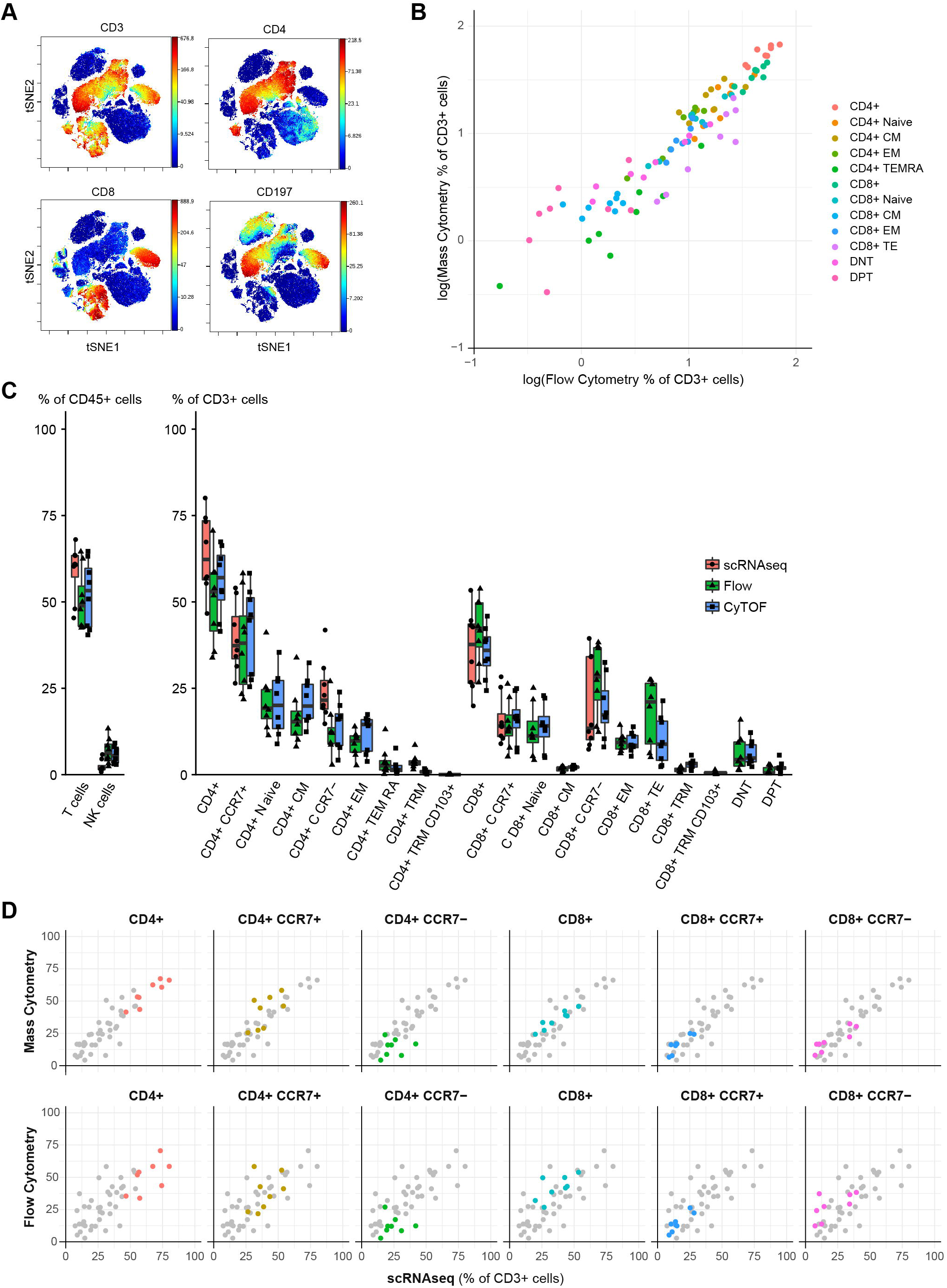
Comparison of single cell RNA sequencing, mass cytometry and flow cytometry assessment of T lymphocyte frequencies in human bone marrow. (A) Mass cytometry for phenotyping of T cell populations visualized using viSNE analysis with expression of key markers shown (B) Comparison of cell frequencies for each donor determined by mass cytometry and flow cytometry. (C) T cell frequencies for cell populations identified by mass cytometry, flow cytometry and single cell RNA sequencing. (D) Individual sample comparisons by scatter plot for each cell population. All population comparisons are shown in background in grey.

To further compare mass cytometry and flow cytometry with scRNAseq of T cell populations, the frequencies of T cell subsets for this cohort of eight donors were determined using all three of these methods, shown in Figure 3C with sample correlations reported in Figure 3D. Comparing frequencies of T cell populations between mass cytometry and scRNAseq confirmed a small but persistent skewing in the identification of NK and T cells. Using Bland-Altman calculations, the mean difference between scRNAseq and mass cytometry for T cells was -6.5% (95% CI: -29% to 16%) and for NK cells was 3.2% (95% CI: -1.1% to 7.6%).

CD8 cytotoxic T cells and NK cells are known to have substantial overlap at the transcriptome level (15). To better understand systemic bias in the frequency of NK or T cells identified, we confirmed that overlapping gene signatures are found in clusters annotated as NK or T cells in this scRNAseq data set (Figure S4). The reasons for this bias are likely however multifactorial.

### Bulk RNA-sequencing

Analysis of bulk sample RNA expression has been used to attempt to deconvolute the proportion of each cell subtype in human tissues (16, 17). Finally, as an additional resource, stranded whole transcriptome sequencing of RNA isolated from thawed BMMCs was performed on samples from all eight subjects for which mass and flow cytometry and single cell RNA sequencing was available. Initial analysis using deconvolution algorithms that attempt to predict the proportion of cell subpopulations is shown in Table S4.

## Discussion

Changes in the immune system (14) and hematopoiesis (18) occur during human aging. Using an unbiased approached based on unsorted human BMMCs, we describe the major cell populations of healthy human bone marrow from a cohort of donors over a wide range of adult age by multiple high-dimensional single cell techniques. This resource serves as a complement to existing data sets that have consisted primarily of younger donors without associated paired immunophenotyping. Our data set provides a resource of scRNAseq, flow cytometry and mass cytometry data for healthy control cohorts across the full range of adulthood providing not only cell population frequencies and characteristics, but also highlighting individual variation in human cohorts.

Using scRNAseq of a total of over 76,000 cells from 20 healthy donors, all the major bone marrow mononuclear populations are identified, and overall population frequencies are comparable to flow cytometry of the matched samples. A primary limitation is distinguishing cell populations such as NK cells and CD8+ effector T cells, which have overlapping transcriptional programs with a small number of distinguishing genes captured by droplet-based scRNAseq. To overcome this limitation and provide additional reference data beyond previous reports of major healthy bone marrow populations by flow cytometry (19) and mass cytometry (20, 21), we used the strength of mass cytometry for high resolution of T cell subpopulations (22), both to validate our flow cytometry results and provide quantification of rare T cell subpopulations within healthy human bone marrow.

As a data resource, these high-dimensional approaches to bone marrow characterization add valuable information on transcriptional and cell surface marker coexpression. The growing number of bioinformatics tools for mass cytometry (23) and scRNAseq (24, 25) will benefit from reference data sets for validation and integrated comparison across techniques. Future opportunities for integrating these data sets include droplet-based sequencing with oligonucleotide-tagged antibodies, including CITE-Seq (26), REAP-Seq (27), and AbSeq (28), which can be compared to this reference set of cell surface protein and transcriptome expression. As techniques (29) and repositories (30) of high-dimensional single cell human data sets are expanded, validating the observed cell identities will be a critical aspect of interpreting large data set analysis.

Additional aliquots of bone marrow aspirate from this cohort together with paired blood samples, that were not yet analyzed, have been stored. Should transformative technologies emerge over the next few years we would be willing, subject to relevant technology transfer and clinical regulatory approvals, to share remaining samples with academic investigators for additional benchmarking and validation. In summary, this resource provides a reference dataset for cell populations in healthy human bone marrow across a wide age range as assessed by multiple single-cell approaches. We show that scRNAseq quantification of marrow-resident cell populations has good concordance with immunophenotyping by flow and mass cytometry with some discrepancies in T and NK subsets. We hope this unique combined dataset will prove useful both to those seeking to refine or innovate bioinformatic algorithms for scRNAseq data and also to those investigators hoping to apply these powerful single-cell technologies in their own research.

## Acknowledgments

This work was supported by the Intramural Research Program of the National Heart, Lung, and Blood Institute of the National Institutes of Health. We appreciate the technical expertise of Brian Sellers and the NIH Center for Human Immunology, Yan Luo, Yuesheng Li and the NHLBI DNA Sequencing Core, Alan Hoofring of NIH Medical Arts and NIH High-Performance Computing. We thank Sheenu Sheela, Blair DeStefano, Janet Valdez and NHLBI research nurses for bone marrow aspirate procedures. The authors would like to thank Neal Young, Cindy Dunbar (both NIH) and Elizabeth Jaffee (Johns Hopkins) for reading and comments, and Efthymia Papalexi and Rahul Satija (both New York Genome Center, NY) for help with scRNAseq analysis and comments.

## Authorship Contributions

KO performed experiments, analyzed data and wrote the manuscript; KL and MG performed flow cytometry experiments and analyzed data; GG and LD analyzed data; PD and PM designed, supervised and analyzed flow cytometry experiments, CL coordinated donor recruitment; CH designed experiments, analyzed data and wrote the manuscript. All authors reviewed the final manuscript.

Figures S1-S4 and Tables S1-S4 can be found in the supplementary information.

